# Insect pests survey of *Prosopis juliflora*, Afar rangeland, Ethiopia

**DOI:** 10.64898/2026.05.04.722396

**Authors:** Belay Beyene Mekonnen, Etay Gittet Lemma, Seid Eshetu Ali

**Author notes:** . Po.box 132.

## Abstract

*Prosopis juliflora* is an invasive alien plant species and a problematic weed that poses significant ecological and socio-economic challenges in Ethiopia, particularly in the Afar rangelands. The study explored the diversity and effects of insect herbivores communities feeding on the flowers and pods of *P. juliflora* to determine their role in limiting reproductive success across three selected ecological sites: Amibara, Gewanne, and Aysayita. A total of 118 adult insect specimens were collected between January and November 2021 using a sweep net and hand collection methods. Community structure, analysis via the Shannon Wiener diversity index, strongly influenced damage pattern. Amibara exhibited the highest insect diversity resulting in significant reproductive damage, including 5.98% of flower loss and 10.39% pods tunneling, primarily caused by Chrysomelidae and Pyralidae. Conversely, Gewanne was showed lower diversity, but higher sap-sucking (13.39 % shriveled pods; 5.11 % flower curling) were caused by Aphididae. Overall, 18.41 % of the pods, and 11.59 % of the flowers were exhibited insect related injury. These finding confirm that more internal seed predation and nutrient depletion were revealed significantly reduce viable seed production. The result was suggested that natural insect communities currently function as partial biological control agents. This indicates strong potential for developing integrated biological control strategies to manage *P. juliflora* invasion in Ethiopia rangelands.

## 1. Introduction

*Prosopis* is a tree native to South America, which was introduced from here to different parts of the world, commonly called mesquite, and is thorny, usually xerophilous shrubs or trees in the family Fabaceae. About 45 species are found worldwide [1].

In Ethiopia, the introduction of invasive alien species (IAS), such as Prosopis (*Prosopis juliflora*), *Parthenium* (*Parthenium hysterophorus*), and Water hyacinth (*Eichhornia crassipes*), has been facilitated by globalisation, international trade, and modern agricultural methods. Around 35 such species now endangered diverse ecosystems across the country, from arid lowland deserts to elevated highland plateaus [2]. *Prosopis juliflora* was introduced in Ethiopia in the late 1970s through collaborative efforts of governments and international development organizations for its adaptability to desert conditions, fast growth, and source of fuel wood, livestock fodder, human food, and bee forage. It also provides shade, stabilizes soil through an extensive root system hence controlling soil erosion, and increases soil fertility through litter and fixing of atmospheric nitrogen as it belongs to the legume family [3, 4]. However, the species rapidly naturalized and expanded into new locations in alarming rate, which neither anticipated nor desired, since its introduction to Ethiopia [5,6], due to the disadvantages and costs of Prosopis for local livelihoods, rangeland health, and biodiversity, and for the national economy because of reduced livestock production family [3]. Using its potential beneficial effects as an opportunity to manage the expansion of this weed in the area where invade (e.g. fuel wood, construction, charcoal production, feeding livestock by crushing pods), chemical treatments (e.g., glyphosate for *E. crassipes*), and biological controls (e.g., *Neochetina weevils*) are the best management option, [2, 3, 4]. However, the approach could not get wider acceptance as there was no immediate benefit to the people, and insisted that Prosopis removed. It is hard and expensive to remove as the plant can regenerate from the roots [4]. [6]; suggest that future intervention in the region should be expanding beyond charcoal production to explore high-value applications, such as human nutrition, medicinal use, and honey production. Therefore, integrate government and NGO efforts to ensure the technology is adopted long-term by the community. In addition, researchers, academicians, and public administrations coordinately should develop a National Strategy and Action Plan aimed at effectively managing the species using a combination of biological, chemical, mechanical, and utilization methods [2, 6].

The resulting decline in pollination rate for native flora significantly undermines local biodiversity and inhibits natural ecosystem regeneration. In addition, the invasion is exasperated by livestock’s when animal consumes Prosopis pods the seeds pass through their digestive tract largely intact. While digestive juice dissolves the outer sugars, they do not compromise the seeds viability. Instead this process facilitates rapid germination within moist feces [4]. Consequently, livestock have become a primary vectors for the dissemination of the *Prosopis* across vast pastoral land, particularly in the Afar Regional State. In response; recent trends, (2024-2025) indicate the Ethiopian management strategy shifting towards Integrated Management. This approach combines mechanical clearing, and charcoal utilization, follow up with fire or chemicals to kill the emerging seedlings, and with the prospective use of biological control agents to suppress reseeding [7, 8]. However, the adverse impact of *P*.*juliflora* expansion on sustainable development of the rangelands of the area and related livestock productivity was higher compare to those in non-invaded areas of Afar region [9]. Therefore, it is critical to study adapted natural insect enemies that target the reproductive structure of this weed to effectively limit its spread.

Given the above four-decade residence time of *Prosopis juliflora* in the study area it is hypothesized that the local entomofauna-particularly those associated with taxonomically related with native Fabaceae have undergone host range expansion. This adaptation likely target the nutrient dense reproductive structure contributed to the observed level of flower and pod infestations or damage, potentially slowing its spread. The primary objective of this study was to identify these “new” natural enemies, assess their abundance and diversity, and quantify the level of damage they inflict on the weed. Such data is essential for determining the feasibility of incorporating these adapted native insects into integrated biological control programs for *Prosopis*.

## 2. Methods and materials

### 2.1. Descriptions of the study area

The studies were conducted at the Lower Awash area (Aysayita) and Middle Awash area (Gewanne and Amibara) area of the Afar region (Fig 1).

**Fig 1.**
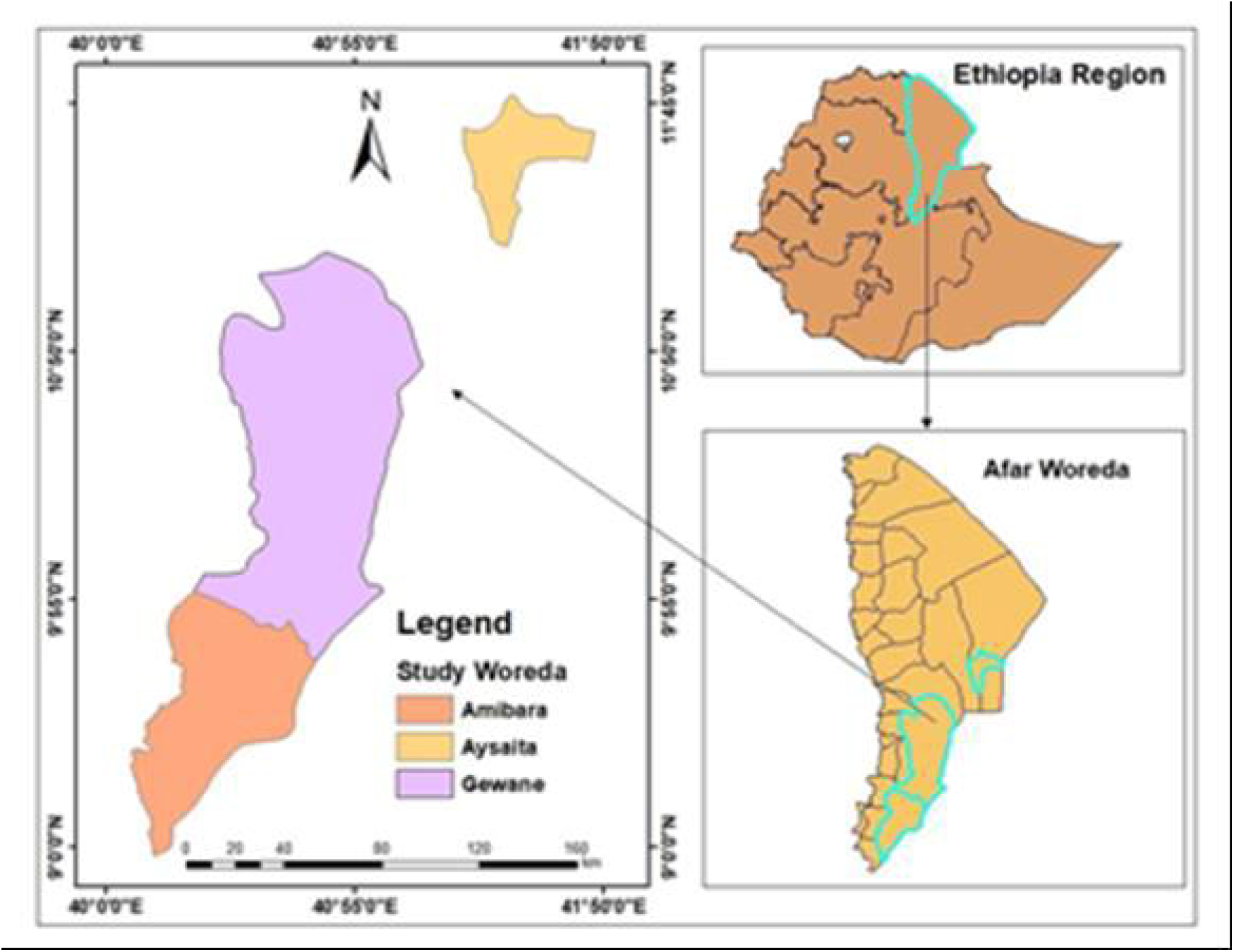
Map showing the study areas (Aysayita, Gewanne, and Amibara) in the Afar Region, Ethiopia. (Samara University GIS Lab 2023)

Aysayita Woredas is situated about 670 km northeast of Addis Ababa. Aysayita town has a latitude and longitude of 11°34′N41°26′E and an elevation of 300 meters (980 ft). The mean annual temperature is between 30 and 45°C, and about 144 mm of the rainfall (Precipitation). Aysayita district is bordered on the south by the Afambo, on the west by the Dubti, then on the north by the Awash River which separates it from the Elidar, and on the east by the Djibouti [10].

The altitude of the middle Awash area (Gewanne and Amibara) ranges from 500 to 820 m above sea level, and it is located between 9° 30′ and 10° 20′ N and 40° 30′ and 40° 50′E. Rainfall in the Awash area ranges from 330 mm to 820 mm. Although extreme temperatures were recorded in the Northern part of Afar, in the central area 35 years of metrological records show that the average lowest temperature is 9 ^0^ C and the average highest temperature is 40^0^ C [11].

### 2.2. Sampling Procedure

Sampling data was conducted every two month intervals from January 2021—November 2021. Sample data collection was done from the roadside of three ecological sites: Aysayita, Amibara, and Gewanne. So, both purposive and simple random sampling methods were used to collect sample insects that adapt and affect the reproductive structures of *P*.*juliflora*. In each study site among twelve line transects two line transects with the size of 100 ms each and with different levels of *P*.*juliflora* invasion were selected purposively. During sampling, three randomly selected quadrates, each measuring 20ms*20ms were chosen for the study.

### 2.3. Data collection

A survey of the insect natural enemies of *P*.*juliflora* was conducted during peak insect activity periods: in the morning (07:00-10:00) and late afternoon (16:00-17:00). Sweep net was used to collect flying insects and hand-collecting was used to sample insect directly from flowers and pods. This timing protocol was implemented because the diurnal activities of most insect is constrained by temperature, with many species becoming inactive or seeking refuge during the extreme midday heat characteristic of the study area to avoid desiccation and thermal stress [12, 13]. Sampling these cooler, more humid periods aligns with known bimodal activity patterns in arid-land insects and increase the probability of collection [14]. Insect specimens encountered in the field were pilot sorted and identified using the available illustrated field guides [15]. Specimens that were difficult on cite were collected as voucher specimens. Adult insects were killed and preserved in vials containing 75% isopropyl alcohol (rubbing alcohol). Consequently, the necessary information such as; sample date, site name, line transects type, and parts of the reproductive structure where the sample specimen was collected was written in code and attached to each vile, and brought to Samara University Biology laboratory. However, to identify immature insects pod-bearing branches were covered with mashed cloth bags. After 15-21 days of incubation period, the bag was recovered, and emerged adults were collected for precise identification [16].

To collect the data for the damage resulting from insect feeding, three randomly selected quadrates, each measuring 20ms*20ms were chosen for the study. From each quadrant three plants were purposely selected for further inspection to study insect communities that inhibit and interact with these plants. Additionally, five branches from each of selected plant, each bearing three to five flowers were collected to examine any signs of damage resulting from insect feeding. Similarly, mature and immature pod sample specimens were gathered from three randomly selected plants of the studied weed. These pods were thoroughly inspected, both externally and internally, for any damage caused by insects, including potential harms of the seeds within. All inspection and potential assessments of insect feeding were conducted in the field as well as in the Biology laboratory at Samara University.

### 2.4. Identification and assessments of the impact of insect pests on reproduction

Identification of insect specimens was carried out based on morphological features using a magnifying lens and binocular microscope. In addition, during identification of the collected insect specimens was supported by the identification guide, insect pictorial keys [15], and related journal articles Furthermore, to assess the impact of insect herbivore on reproduction, collected *P. juliflora* pods were inspected externally and dissected internally insides. Any damage symptoms (tunneling of the pod, internal galleries, fras or seed consumptions) were precisely recorded, following established protocol for assessing pre-dispersal seed predation [17].

### 2.5. Data analysis

Data analyses of insect diversity were measured in the study area. An Excel data sheet was used to record and organize the collected data. Shannon – Wiener diversity index was used to measure the diversity of insect families among the study sites [18].

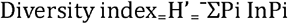

Where Pi = S / N, S = the number of individuals of one species. N = the total number of all individuals in the sample and In = logarithm to base e. A species with a higher value of H’ is more diverse than species with a lower value of H’.

Evenness (E) is an index that makes the H’ values comparable between communities by controlling for the number of species found within the communities. For a given number of species in a community, the highest H’ (H’max) represented as:

H’max_=_ lnS (Where S = the total number of species). So, when H’ (actual estimate of the community) is divided by H’max, is Evenness.

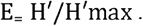

Here, (E) can range from close to 0, where most species are rare and just a few are abundant, to 1, where the potential evenness between species (H’max) is equal to that which will be observed (H’). This Means insect with larger E value has more even distribution Patterns of relative abundance of species determine the dominance component of diversity. In this study, the relative dominance of each family within sit was determined by calculating the dominance index using the following formula:

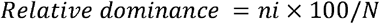

Where ni = the number of Phytophagous insects in the ‘i’ ^th^ family, and N = the total number of Phytophagous insects in all the families collected in each site.

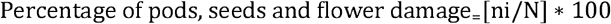

Where ni = the number of damaged pods, seeds or flowers in the ‘i’ ^th^ of damage type, and N = the total number of pods, seeds or flowers collected in each site (the total number of pods, seeds or flowers collected in the study area).

## 3. Results

### 3.1. The diversity and abundance of insect pest communities on *P. juliflora* in the study area

In Ethiopia, *Prosopis juliflora* belongs to the first invasive weed [3]. According to [19], the invasion rate is estimated at 50,000 hectares per year, in Afar Regional Administration. Considering the above information, the expansion nature of the weed and the livestock-mobile nature of the area enable us to guess its invasion coverage of more than two million in now a day. Thus, to restrict such an alarming expansion rate of the weed would be defensible by combining more than one control option.

Therefore, the objective of this study was to identify insect natural enemies that adapt to feed and affect the floral, pod, and seeds, including the abundance and diversity of these pests in the study area. Consequently, 233 insect individuals were collected from January 2021— November 2021. All of them were identified at family levels, 132 individuals at genus levels and 29 individuals of them were identified at the species level (Appendix 1).

Among the collected insect communities, 118 of them were identified as insect pests that adapted to feed and affect the reproductive part of *P. juliflora*. These insect pests were belonging to 4 orders and 7 families (Table 1). The remaining 115 insect individuals were identified as non-reproductive part feeders, pollinators, and predators (Appendix 1).

**Table 1.** Family composition and ecological impacts of insect herbivores associated with *Prosopis juliflora* reproductive structures across three study sites of the Afar region, Ethiopia.

| Family |  |  |  | Feeding Guild and Ecological Impacts |
| --- | --- | --- | --- | --- |
|  | Amibara | Gewanne | Aysayita |  |
|  | (n=50)<br>n (%) | (n=45)<br>n(%) | (n=23)<br>n(%) |  |
| **Meloidae** | 4(8.0) | 2(4.44) | 3(13.04) | Feed on flowers and pollen, preventing Pollination [20]. |
| **Chrysomelidae** | 3(6.0) | 2(4.44) | 0(0.00) | Larvae develop inside the seeds destroying them entirely [21]. |
| **Psyllidae** | 4 (8.0) | 3(6.67) | 2(8.7) | Sap-suckers that attack flowers racemes and young pods [22]. |
| **Aphididae** | 19(38.0) | 23(51.11) | 10(43.48) | Sap-suckers that attack flowers racemes and young pods [23]. |
| **Lygaeidae** | 3(6.0) | 3(6.67) | 1(4.44) | Pierce the pod to suck nutrients from developing embryo [21]. |
| **Acrididae** | 13(26.0) | 9(20.0) | 7(30.43) | Generalist that frequently consume flowers and succulent green pods [20]. |
| **Pyrallidae** | 4(8.0) | 3(6.67) | 0(0.00) | Borers that tunnel through pods and consume the internal pulp and seeds [16, 21]. |
| **Total** | 50(100) | 45(100) | 23(100) |  |

Table 2 was showed the dominance, diversity, and evenness indices of insect pest families associated with the reproductive part of *P. juliflora* for the study sites: Amibara, Gewanne, and Aysayita. Consequently, the Amibara was remains the site with the highest diversity (*H*′=1.662) and evenness (*E*=0.854) for the reproductive pests community. However, Aphididae and Acrididae emerged as the most frequent taxa across all sites, together accounting for more than half of the total individuals were collected (Table 2).

**Table 2.** Diversity indices of insect herbivore families associated with the reproductive structures of *Prosopis juliflora* across three study sites of the Afar region, Ethiopia.

| Site | Total<br>individuals(N) | Family<br>Richness<br>(S) | Dominance<br>Index (D) | Diversity<br>Index ($H'$) | Evenness<br>index<br>(E) |
| --- | --- | --- | --- | --- | --- |
| **Amibara** | 50 | 7 | 0.2384 | 1.662 | 0.854 |
| **Gewanne** | 45 | 7 | 0.3185 | 1.483 | 0.762 |
| **Aysayita** | 23 | 5 | 0.3081 | 1.339 | 0.832 |

As far as, Gewanne was showed a lower evenness (*E*=0.762) compared to Amibara (Table 2). This was largely driven by the high dominance of Aphididae (was made up over 50% of the reproductive-part feeders at this site) (Table 1). This suggests that in Gewanne, sap-sucking pressure on flowers and young pods was the primary reproductive constraint. On the other hand, Aysayita had fewer families (S=5) due to absence of Chrysomelidae and Pyralidae, reducing overall diversity despite moderate evenness (E=0.832) compared to Gewanne (Table 2). These differences likely reflect site-specific ecological factors affecting pest family composition.

Overall, by focusing purely on these families, the data suggests that Amibara study site was faced with the most “diverse” attack on its reproductive system (multiple different ways the plant is hindered), whereas Gewanne was under heavy, specialized pressure from sap-sucking insects (Aphididae), which likely leads to significant flower loss before pods can even form.

### 3.2. Damage of pods and flower observed during the study

During the sampling period, the sign of damage of pods from collected pod samples of this weed were observed are; tunnels and holes, shriveled or deformed, abnormal swollen and gal formation, and tiny puncture mark or scar (Table 3.). Likewise, the sign of flowers damages were observed are; Honey dew & curling of flower petals, wilting flower, and Loss of flowers (Table 4.). The aforementioned damaging types of Prosopis pods and flowers were observed from the result of the left over by the feeding of insects at some stage of their life cycle. Hence, such symptoms were revealed an adaptation of some insect pests that feed pods and flowers, which causes damage to this weed.

**Table 3.** Percentages of Prosopis Juliflora pod damage by category across three ecological sites in the Afar region, Ethiopia.

| Study site | n<br>Pods | Tunneling/Holes<br>n (%) | Shriveled/Deformed<br>n (%) | Punctures/Scars<br>n (%) | Total<br>damage<br>n (%) |
| --- | --- | --- | --- | --- | --- |
| Aysayita | 240 | 0(0.00) | 0(0.00) | 9 (3.75) | 9(1.27) |
| Gewanne | 235 | 11(4.68) | 31(13.39) | 20 (8.51) | 62(8.87) |
| Amibara | 231 | 24(10.39) | 23(9.96) | 12 (5.19) | 59(8.36) |
| **Total** | 706 | 35(4.96) | 54(7.65) | 41(5.81) | 130(18.4) |

**Table 4.** Percentages of *Prosopis Juliflora* flower damage by category across three ecological sites in the Afar region, Ethiopia.

| Study site | n samples | Honeydew/Curling<br>n (%) | Wilting<br>n (%) | Flower loss<br>n (%) | Total damage<br>n (%) |
| --- | --- | --- | --- | --- | --- |
| Aysayita | 192 | 11( 5.73 ) | 1(0.52) | 3 ( 1.56 ) | 15( 2.71 ) |
| Gewanne | 176 | 9 (5.11) | 6 (3.4 ) | 10 ( 5.68 ) | 25( 4.53 ) |
| Amibara | 184 | 6 (3.26) | 4 (2.17) | 11( 5.98 ) | 21( 3.74 ) |
| **Total** | 552 | 26 (4.72) | 11(1.99) | 27( 4.89 ) | 64( 11.59 ) |

Consequently, during the study season 1402 pods were collected. These collected pods 480 were from Aysayita 470 from Gewanne, and 462 from Amibara study sites. The number and percentage of damages were computed within the study sites according to the damaging type were observed. The highest percentage of pod damage was observed at the Gewanne 8.78 %, followed by Amibara 8.36 % with the least damage recorded at Aysayita and the least was 1.27%. Finally, percentage the pods which show symptoms of damages by insect pests were computed with in the study area, 18 % of the pods were damaged (Table 3.).

The damage symptom shown in the table 3, highly diagnostics of specific “pod specialist” insects such as: the percentage of the tunnels and holes of the pod at Amibara study site 10.8% (Table 3). This was the sign of the Chrysomelidae and Pyralidae larvae (grub/ caterpillar) feed inside the pod; when they reach adulthood, they chew “exit holes” to emerge. Amibara, having the highest diversity (H′=1.662), shows the most severe “tunneling” damage, likely due to a combination of these two families (Table 1 and 2). On the other hand, the highest percentage of the damage symptom shriveled or deformed of pods was observed at Gewanne study site (13.39%) (Table 3), this damage was primarily caused by Aphididae and Psyllidae. Because they suck large amounts of sap during the early “green pod” stage, the pod loses turgor pressure and fails to develop fully, resulting in a shriveled appearance. The high Aphid dominance in Gewanne over 50% (Table 1), was correlated perfectly with this being the site’s most frequent damage type. In addition, the pod damage symptoms of the tiny puncture mark or scar at Gewanne also were recorded the highest 8.51% (Table 3). These were “stings” from Lygaeidae (Seed Bugs). These bugs use needle-like mouthparts to probe through the pod wall to reach the seed embryo. Each probe leaves a tiny necrotic scar or puncture mark (Table 1).

Table 4, shows the types of flower damage and the extent of damage caused by insect pests on *P. juliflora* in the study sites. The highest percentage of flower damage was observed at the Gewanne 4.53%, followed by Amibara 3.74%, with the lowest damage was recorded at Aysayita and the least was 2.71%. Overall, the percentage of flower inflorescences damaged by insect pests on the study area was 11.05% (Table 4).

As far as the association of insect pest of the studied plant and flower damage was concerned, in Aysayita the dominant reproductive pests were Aphididae and Acrididae (Table 1), the key damage observed was honeydew/curling (5.73%) (Table 4). While flower loss was lower here, the curling suggests a persistent presence of sap-suckers early in the flowering stage, Aphididae was the cause visible curling. However, in Gewanne the key flower damage observed was high wilting and loss (Table 4), and had the highest Aphid concentration (Table 1). This matches the high “Total damage” (4.53%) and significant wilting (3.4%), as heavy Aphid colonies drain the flower’s energy.

Amibara was observed the highest flower loss (5.98 %) (Table 4), and Show high diversity of all families (Table 2). Because Amibara has the highest diversity (H′=1.662), the flowers were being attacked by multiple groups (beetles, bugs, and grasshoppers). This “multi-front” attack results in the highest rate of total flower loss.

## 4. Discussion

The expansion of *Prosopis juliflora* in the study area is mainly the livestock that feeds on pods for their nutritional requirement. However, the indigestible nature of the seed coat of this weed makes it able to pass across the digestive organs of herbivore animals without damage. Moreover, the undamaged seeds of *P. juliflora* drop along with the feces of these animals [3].Consequently, the weeds seeds disseminate over most of this arid area following the movement of the livestock. That is why the question of this study was whether some herbivore insects adapted and affect the flowers, the pods, and the seeds of these weeds. Therefore, understanding the diversity and abundance of insect natural enemies of this weed that cause the damage of the reproductive parts help to find an additional controlling option as a biological controlling agent in the study area. Similarly [24]; mentioned in their study, Pre-dispersal seed predation by insects is a major select force modifying many aspects of seed biology. Insect infestation produces seed or fruit abortion, or both, as, affects seed germination, and modifies fruit color, size, and shape, consequently influencing seed dispersal by animals.

The study revealed a significant association between insect diversity indices (*H*′) and the intensity of reproductive damage across the three sites. Amibara, which exhibited the highest diversity of reproductive pests (*H*′=1.662; *E*=0.852). The distribution of pod damage across the study sites reflects the specialization of the resident insect communities. The high incidence of ‘tunnels and holes’ in Amibara (10.39%) corresponds with the high diversity of internal feeders such as Chrysomelidae and Pyralidae. As noted by [21]; such internal feeding directly compromises seed integrity.

Conversely, the high rate of ‘shriveled or deformed’ pods in Gewanne (13.39%) is likely a physiological response to the dominant Aphididae population. These sap-suckers deplete the pods of vital nutrients during the elongation phase, a phenomenon documented in other legumes by [16]. Amibara also recorded the highest total flower loss (5.98%). This suggests that a more diverse pest complex—comprising flower-feeders like Meloidae, pod-borers like Pyralidae, and generalists like Acrididae—exerts a multi-faceted pressure on the plant’s reproductive potential. In contrast, Gewanne showed the highest levels of wilting (3.4%) and honeydew-related curling (5.11%), which aligns with the dominance of Aphididae at that site. As noted by [22, 25]; sap-sucking insects on *Prosopis* species are primary drivers of flower abortion (drop) due to the depletion of phloem nutrients and the injection of toxic saliva.

The damage observed in this study mirrors patterns found in other members of the Fabaceae family. Scholars like [26,27]; have documented that Meloidae (blister beetles) are gregarious feeders that can destroy over 80% of flowers in leguminous crops such as *Baptisia australis* and pearl millet, significantly reducing pollination success. Similarly, the high damage to pods and seeds recorded across the sites, particularly the internal boring, is consistent with the behavior of Chrysomelidae. Research work by [28, 29]; emphasizes that seed beetles are specialized predators of desert legumes; their larvae develop internally, rendering seeds non-viable and directly limiting the recruitment of new seedlings. These findings suggest that the insect complex at the study sites is not only affecting the immediate seed crop of *P. juliflora* but is utilizing the plant’s reproductive structures in a manner typical of major legume pests globally.

In addition, 18.41 % of the collected pods samples, and 11.59% flowers inflorescences were shown the different damaging symptoms due to the result of the left over by the feeding of insects at some stage of their life cycle. These were the significant factor in limiting the reproductive potential of *P. juliflora*. Because of direct Consumers of the insect families like Meloidae and Acrididae were likely responsible for the physical loss of pods and flowers through direct consumption. And also, the physiological stressors caused by individual insects pests that belong to the family Aphididae and Psyllidae are the drivers of the “Honeydew” and “Wilting” symptoms, which indirectly prevent pod formation by weakening the flower.

The data indicates that the reproductive health of *P. juliflora* in the Afar region was subjected to two distinct types of ecological pressure. These were the internal predation focusing to the Amibara study site was revealed the prevalence of tunnels and holes in this study site due to a high population of seed beetles (Chrysomelidae) and Pyralidae moths. These insects are “seed predators.” Their impact was the most severe because they consume the embryo, rendering the seed non-viable so therefore directly reduces the plant’s ability to spread via seed [16, 21]. On the other hand, the physiological stress focusing to the Gewanne study site was revealed the high percentage of shriveled pods and puncture marks in Gewanne points to Hemipteran (sap-sucking) dominance. While these insects may not always “kill” the seed, they reduce the pod’s biomass and can lead to “empty” or underdeveloped pods. As noted by [29]; such damage affects the nutritional quality and germination vigor of the remaining seeds.

Overall, the diversity and abundance of the insect pest communities on the reproductive part of *P. juliflora* show there are insect natural enemies that were adapted to feed and affect this weed. Therefore, the study will be encouraged to exploit and evaluate the potential use of the insect natural enemies as a biological control agent for the *P. juliflora* in Ethiopia including the Afar region.

## 5. Conclusion

The findings of this study provide a complete assessment of the entomological factors limiting the reproductive success of *Prosopis juliflora* in the Afar region. The results demonstrate a clear link between insect family diversity and the specific types of damage observed on the flowers and pods of the host plant across the three study sites.

A survey study of natural enemies of insects that feed and affect the reproductive part of *P. juliflora* reveals there were a significant association exists between high species diversity and cumulative reproductive impact. Amibara, which recorded the highest diversity index (*H*′=1.66), experienced the most severe mechanical damage, specifically flower loss (5.98%) and pod tunneling (10.39%). This indicates that a diverse pest guild creates a “multi-front” pressure that is more damaging than the dominance of a single species.

In contrast, Gewanne exhibited a lower diversity index but higher specific damage in the form of shriveled pods (13.39%) and flower curling (5.11%). This is attributed to the overwhelming dominance of Aphididae, suggesting that in certain environments, specialized sap-sucking pressure can be the primary constraint on fruit development.

The presence of Chrysomelidae (seed eaters) and Pyralidae in Amibara and Gewanne ecological sites represents the most critical threat to the plant’s invasiveness. By tunneling into pods and consuming seed embryos, these insects act as direct seed predators, effectively reducing the soil seed bank and the long-term recruitment potential of the species.

The damage patterns recorded—ranging from the physiological wilting caused by Hemipterans to the destructive feeding of Coleopterans—is consistent with established scholarship on the Fabaceae family. This confirms that the native insect complex in Ethiopia has successfully adapted to utilize *P. juliflora* as a primary reproductive resource.

Given the high rates of seed and pod damage (reaching an overall total of 18.41% across sites), future management strategies should consider these insect families as potential components of an Integrated Pest Management (IPM) or biological control framework. Specifically, the role of seed beetles should be further investigated for their potential to limit the further spread of this invasive species in Ethiopia.

## Acknowledgment

Primarily, we would like to thank the department of biology access to the biology laboratory. This enabled us to carry out our research with the necessary resources and equipment. We would also like to extend our appreciation to Aysayita, Amibara, and Gewanne Agro-pastoral and Pastoral bureau for allowing us to conduct our research on the Prosopis site. Your willingness to provide us with access to this site was invaluable to the success of our project. Furthermore, we would like to acknowledge the extension workers of the above-mentioned bureau for their support during data collection. Their assistance and guidance were essential to the accurate collection and analysis of our data. Finally yet importantly, we would like to express our gratitude to all those who assisted us in various capacities during the course of our project. Once again, thank you for your support and cooperation. We could not have achieved our research goals without your help.

## Appendices

**Appendix 1:**
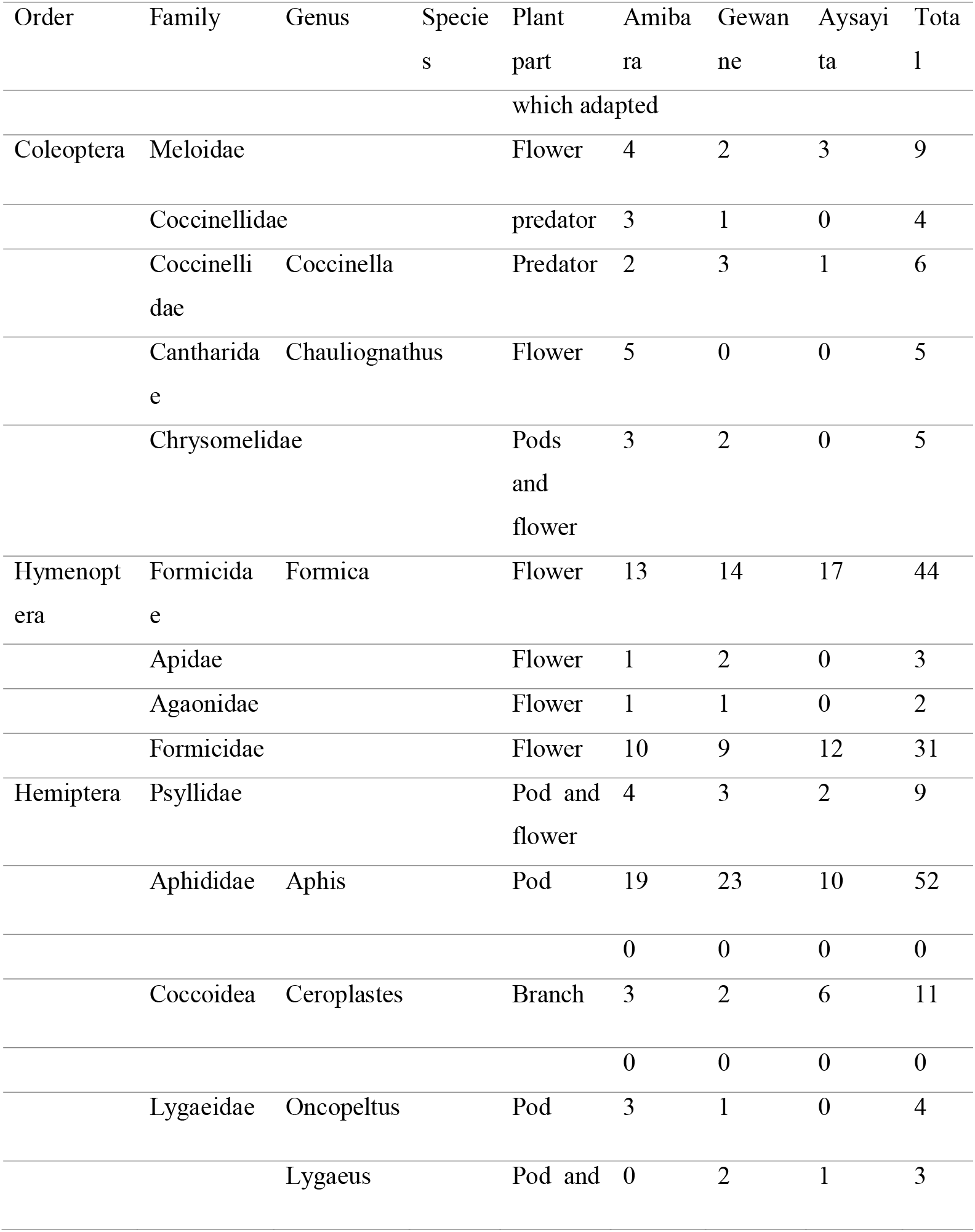
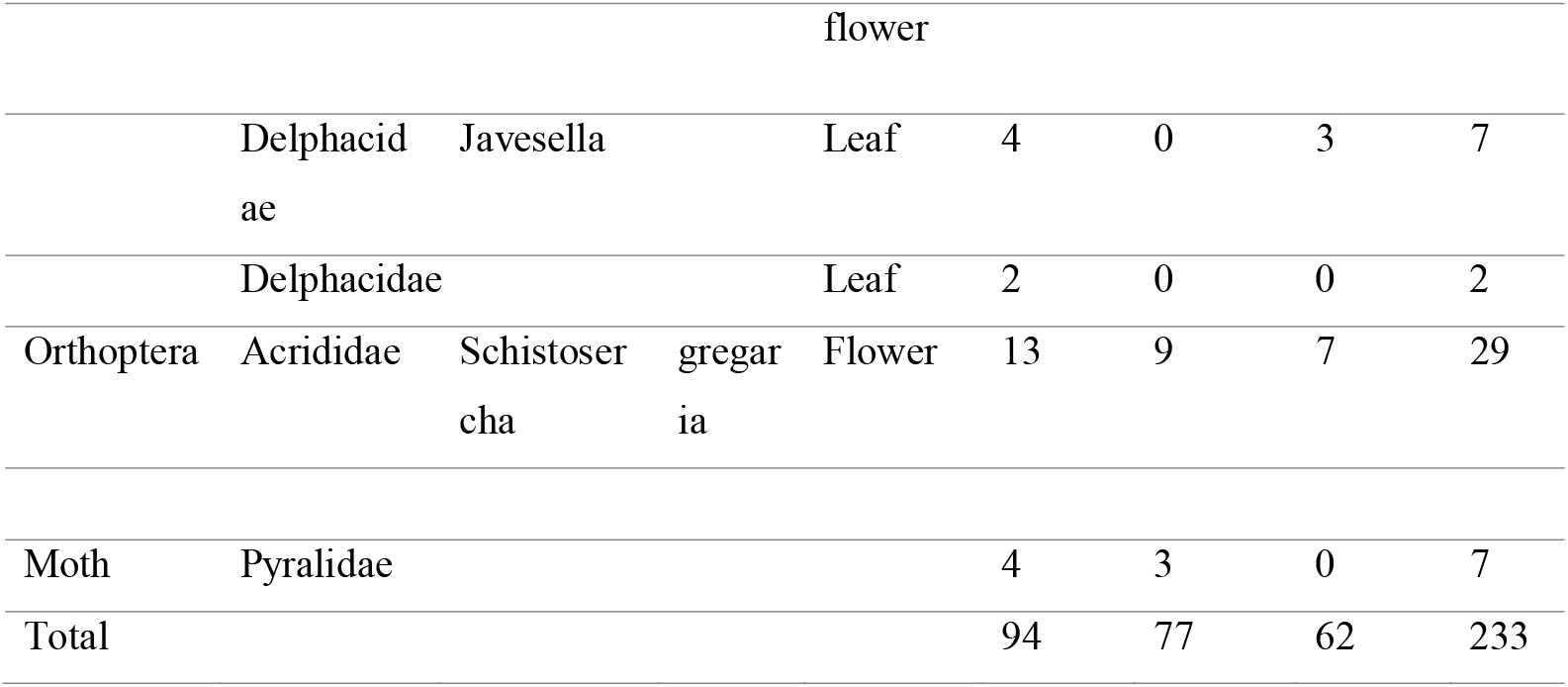
List of insect communities were collected and identified from Prosopis Juliflora across the three ecological sites of Afar region, Ethiopia.

